# Seasonal eDNA-based monitoring of *Batrachochytrium dendrobatidis* and amphibian species in Norway

**DOI:** 10.1101/2022.08.02.502273

**Authors:** Omneya Ahmed, Johan Andersson, Pedro M. Martin-Sanchez, Alexander Eiler

## Abstract

Freshwaters represent the most threatened environments with regard to biodiversity loss and therefore there is a need for national monitoring programs to effectively document species distribution and evaluate potential risks for vulnerable species. The monitoring of species for effective management practices is, however, challenged by insufficient data acquisition when using traditional methods. Here we present the application of environmental DNA (eDNA) metabarcoding of amphibians in combination with quantitative PCR assays for an invasive pathogenic chytrid species (*Batrachochytrium dendrobatidis-Bd*), a potential threat to endemic and endangered amphibian species. Statistical comparison of amphibian species detection using either traditional or eDNA-based approaches showed weak correspondence. By tracking the distribution of *Bd* over three years, we concluded that the risk for amphibian extinction is low since *Bd* was only detected at five sites where multiple amphibians were present over the sampled years. Our results show that eDNA-based detection can be used for simultaneous monitoring of amphibian diversity and the presence of amphibian pathogens at the national level in order to assess potential species extinction risks and establish effective management practices. As such our study represents suggestions for a national monitoring program based on eDNA.

## Introduction

Freshwater environments are threatened by biodiversity loss, and there are over 100 documented cases of extinction in such environments during the 2000s (Dudgeon et al. 2006; Tickner et al. 2020). In the case of amphibians, 30% of all freshwater amphibians species are threatened by extinction (Dueñas et al. 2021). The spread of invasive species because of human activity has been proposed to be one of the greatest threats to indigenous species (Beaury *et al*. 2020). One such invasive species is the chytrid fungus *Batrachochytrium dendrobatidis* (*Bd*) which causes the disease chytridiomycosis in amphibians. Chytridiomycosis has led to large reductions in the populations of amphibian species, and in some cases extinction across the world (Berger et al. 1998; Laurance et al. 1996; Longcore et al. 1999; Mendelson et al. 2006).

Tickner et al. (2020) recently presented an emergency recovery plan for freshwater biodiversity, which includes protecting and restoring critical habitats, and preventing the introduction and spread of non-native species. In addition international treaties such as the global biodiversity framework under the Convention on Biological Diversity and the EU’s biodiversity strategy for 2030 have been initiated. The later has been established to facilitate an EU-wide restoration plan and network of protected areas, as well as introducing measures to enable necessary transformative changes and to tackle the global biodiversity challenge. Here monitoring programs of biodiversity are a prerequisite to design, implement and validate these international conservation and restoration efforts.

To be successful, biodiversity monitoring must be designed in such a way as to minimize the main sources of error (Skalski and Robson, 1992; Thompson et al. 1998). A common source of error comes from the fact that all survey methods fail to detect all individual species in an ecosystem (Mathieu et al. 2020). A second source of error arises from the difficulty in efficiently investigating large areas, meaning that conclusions must be based on a low number of samples taken from a few locations. This is compounded by the fact that many collection strategies are based on subjective assessments of how representative certain sampling locations are, or how easily accessible sites are. Thirdly, environmental managers face major problems in identifying and counting species, since organisms can be difficult to detect or to distinguish from one another. Hence, there is a need for novel methods to survey biodiversity at the national levels.

Technological advances over the past decades in molecular biology now allow us to minimize some sources of error. Environmental DNA (eDNA) analysis is a rapidly evolving methodology used for studies of current and past biodiversity (Valentini et al. 2009; Taberlet et al. 2012). eDNA has broad applications in: the analysis of biodiversity in microbes, plants, and animals (Eiler and Bertilsson, 2004; Zinger et al. 2012; Valentini et al. 2016), analysis of diet (Deagle et al. 2005; Pompanon et al. 2012), reconstruction of past biodiversity or environmental changes (Jørgensen 2012; Gigue-Covex et al. 2014; Langenheder et al. 2016; Parducci et al. 2013), and environmental monitoring (Jerde et al. 2011; Eiler et al. 2013). In principle, biodiversity across the entire tree of life present in a particular system can be assessed by DNA metabarcoding (Stat et al. 2017). Substantial advantages of eDNA-based methods are higher cost and time effectiveness compared to many traditional survey methods (Evans et al. 2017), their noninvasive nature (Cristescu & Hebert 2018), and high specificity and sensitivity (Wilcox et al. 2013).

Testing for the presence and concentration of microbial pathogens such as *Bd* using eDNA methods is appealing because it relies on noninvasive sampling and free-living stages persistent over several weeks can be detected (Brannelly et al. 2020). Similarly, amphibian diversity can be assessed without the need of finding and capturing animals in the environment, thus avoiding harming animals and introducing sampling biases (Valentini et al. 2016). Despite the high sensitivity of eDNA methods, species detection by eDNA approaches is not free from sources of error similar to those found using conventional methods. In fact, incomplete detection is inevitable when assessing the presence/absence and number of species, regardless of the methods used (MacKenzie et al. 2006). This can be partially overcome by the high sensitivity and capacity of eDNA methods that allow more samples to be processed leading to a higher likelihood of detection, than when conventional methods are used. In addition, adaptive sampling can increase the likelihood of detection; for example, by sampling during the optimal season and adapting spatial sampling strategies to the target organisms (Buxton et al. 2018).

In this study, we aimed to assess the distribution of the chytrid fungus *Bd* from water samples by employing eDNA methods. *Bd* is regarded as a generalist fungal pathogen, presently known to occur in Sweden (Rosquist 2020) and Denmark (Scalera et al. 2008), but little is known about the spread and virulence of *Bd* in Norway (Taugbøl et al. 2021). Two qPCR assays and a metabarcoding approach were used to detected *Bd* and amphibian species detection, respectively, throughout Southern Norway. The specific objectives of this study were to (i) assess the spread of *Bd* in the southeastern part of Norway over several years, and (ii) explore potential risks for amphibian hosts using co-occurrence analysis and reviewing the literature. Lastly, (iii) we also intend to provide methodological suggestions for eDNA based national monitoring programs.

## Materials and Methods

### Sampling design

Sampling sites located in Southern Norway (regions of Viken, Oslo, Vestfold/Telemark, Agder and Innlandet) were chosen randomly from sites with observations of amphibian species as reported in the Norwegian species database “Artsdatabanken”. Resampling for each month was done on the complete database, thus facilitating that sites were sampled multiple times. As the density of sites with species observations is related to human presence, our sampling focused on anthropogenic impacted and potential *Bd* positive sites. Introduction of Bd is proposed to be related to human activities (VKM report). During 2019, 110 bodies of water, including forest-, agricultural- and urban ponds, were sampled. To test for the optimal sampling season, we collected and analyzed samples from three different times in 2019: May or early-June (37 samples), July (36 samples) and September (39 samples). These sampling times correspond to life history stages such as spawning, post-spawning and juveniles, respectively (See supplementary Table S1). To compare the detection probabilities between the traditional and eDNA-based sampling, 91 bodies including forest, agricultural and urban ponds, and 10 swabs from various amphibian specimens were collected over a single season in 2020. Swabs were taken from frog and salamander skins at Hasseldalen (1 swab), Værmyr (3 swabs), Ringveien 52 (3 swabs) sites, while one swab from captured tadpoles were collected from Butjenna, Øyenkilen, Sodra Skauen Gård waterbodies (See supplementary Table S2). Swabs and tadpoles were preserved in 70% ethanol on site and stored at 4 °C until further analysis. Details about the survey design including geographical and temporal distribution of the samples are given in supplementary Figure S1.

Water samples were collected with a beaker attached to expandable sampling pole (a device that can reach 4 meters). At least 5 samples (of 20-100 ml) per site were pooled together. We used cartridge filters (Sterivex filter units 0.22 μm pore diameter, Millipore, Merk, Darmstadt, Germany) and sterile 50-ml syringes to filter between 75 (minimum) and 1440 ml (maximum) of pooled water samples from each site (median volume of 480 ml). Sampling was performed by the same two persons throughout the study. A syringe - cartridge filter setup allowed for reproducible sampling without the need of heavy equipment and access to electricity. Filters were immediately frozen in dry shippers cooled with liquid nitrogen and then stored at −80°C in the laboratory until further analysis. Chemical and physical parameters such as temperature, conductivity, pH, and turbidity were also measured on site (See supplementary Tables S1 and S2).

### DNA extraction and *Bd* qPCRs

All lab benches and equipment were cleaned by using 5% sodium hypochlorite and 70% ethanol in the field and laboratory. This included the three laboratories used for DNA extraction, prePCR or postPCR. Here pipettes and consumables were subjected to UV exposure for 30 min prior to usage. For water samples, DNA was extracted from the sterivex filters using a Qiagen DNeasy PowerWater Sterivex Kit (Qiagen, Germany) including one unused filter as a negative control in one of the DNA extraction runs. The collected swabs were removed from ethanol and left to dry in clean Eppendorf tubes for 2 hours at 50°C to evaporate residual ethanol. The DNA extraction was subsequently performed using the Qiagen Blood & Tissue DNA extraction kit. The quantity and quality (280/260 ratio) of the extracted DNA were assessed using NanoDrop.

### *Bd* qPCR assays

We used two TaqMan assays amplifying different regions of the ribosomal RNA operon (ITS1 – 5.8S rRNA gene – ITS2). First the Boyle TaqMan assay (Boyle et al. 2004), targeting the Internal Transcribed Spacer 1 (ITS1) region where qPCR mixture contained 10 μL of 2X BioRad SsoAdvanced Universal Probes Supermix, 0.8 μM of the forward primer ITS1-3 Chytr (5’-CCTTGATATAATACAGTGTGCCATATGTC-3’) and the reverse primer 5.8S Chytr (5’-AGCCAAGAGATCCGTTGTCAAA-3’) and 0.24 μM of Chytr MGB2 (FAM-5’ TTCGGGACGACCC-3’-NFQ-MGB), 1 μL of internal extraction control (IPC, if not co-extracted with the sample), 1 μL IPC primer and probe mixture, 5 μL of environmental DNA sample and RNase/DNase free water to reach 25 μL as final volume. The second assay was based on the commercially available product *Batrachochytrium dendrobatidis* 5.8S rRNA Advanced Genesig Kit (Primer design Ltd, UK) which was carried out according to manufacturer’s instructions. Both qPCR assays were conducted in a BioRad CFX96 thermocycler (USA) using the following program 2 min at 95°C, followed by 50 cycles of 10 s at 95°C and 1 min at 60°C. At least 8 serial standard dilutions were tested and validated before choosing five standard dilutions (1000, 100, 10, 1 and 0.1) to perform qPCR runs with environmental samples in triplicates. *Bd* positive controls were added in post PCR room while the qPCR master mix was prepared in pre-PCR room.

For obtaining the standard curves needed to quantify *Bd* in environmental samples with the Boyle TaqMan assay, we used a DNA extract from *Bd* spores provided from Professor Anssi Laurila’s research group at Uppsala University. This extract had an expected concentration of 100 genomes (or genome equivalent) per µL. The number of rRNA operons in each spore was not known therefore concentration was given as genome equivalent. Samples collected in 2019 was analyzed with only the commercial assay while samples collected in 2020 were analyzed by both assays in triplicates including two negative controls in 96 well plates. Positive Bd sample was estimated if 3 or 2 of the three qPCR runs generated a positive Bd signal within the range of serial dilutions of the standard. If only 1 of the three qPCR runs was positive an additional fourth qPCR run was conducted. Sanger sequencing was analysed to confirm the amplification of the target fragment with Ct value > 39.5. To compare our results with known occurrences of Bd, we included data from a previous Norwegian survey in 2017 (Taugbøl et al. 2021).

In addition, to further assess the sensitivity of this assay and estimate the target copy numbers (ITS1 copies), we constructed an extra standard curve using a synthetic gBlocks double-strain ITS1 fragment (“Bd_26-271”, this name refers to the positions in the reference sequence NR_119535; 246 bp) based on eight decimal serial dilutions from 0.39 to 3.9 x10^6^ ITS1 copies per reaction. The limit of detection (LOD) was estimated using a discrete approach and defined by the highest dilution where at least a single replicate of the triplicated standard dilution of 5 serial concentrations showed amplification in the respective assay consistently throughout all qPCR experiments with at least 95% confidence (modified from Klymus et al. 2019). To define the limit of quantification (LOQ), we identified the highest dilution where a R^2^ > 0.98 and an efficiency between 85 and 115% was obtained when fitting a linear regression to log-transformed *Bd* quantity and Cq values of the dilution series for each individual experimental (qPCR) run.

### Internal control

In order to determine the DNA extraction efficiency and the presence of inhibitors, a separate internal extraction control DNA, primers and probe mixture with different target region, IPC, was provided with the *Batrachochytrium dendrobatidis* 5.8S rRNA Advanced Genesig Primer design kit (United Kingdom). In 2019, the batch of samples collected in September were co-extracted with IPC by mixing 4 μL of IPC with Qiagen lysis buffer (as described in the manufacturer’s instructions) and tested in a qPCR run. In 2020, 30 randomly selected DNA samples were co-extracted with IPC while the remaining extracted DNA samples were spiked with the IPC. To spike DNA with the internal control, the internal control was diluted 1:20 and 1 μL was added to a subset of samples which did not undergo internal control co-extraction, prior to the qPCR. The internal control is detected through the VIC fluorophore channel. We assumed a 100% extraction efficiency if Cq value of 27 was observed with Cq values of 27±2 within the normal range. Samples with Cq values of 30 or higher were considered to have PCR inhibitors.

### Amphibian mock community

Mock communities are now widely used in metabarcoding as a positive control and to validate analysis procedures from library preparation to bioinformatics analyses. A mock community of five amphibian species was prepared from samples provided by the lab of Anssi Laurila (Uppsala University) and Nordens Ark, including DNA extracts of *Rana arvalis* and *Bufo bufo,* and swabs provided by Anssi Lab (preserved in 95% ethanol) of *Bufotes variabilis*, *Epidalea calamita,* and *Pelophylax lessonae*. DNA extraction from the swabs was conducted as described above. Quantification of the DNA concentration was performed using the PicoGreen assay (ThermoFisher, US). Approximately 10 ng of the extracted DNA from each species were mixed in equal amounts and diluted to prepare 5 dilutions: 1.0, 0.5, 0.1, 0.01 and 0.001 ng/μL.

### Amphibian DNA metabarcoding-Library preparation

Paired-end sequencing on the Illumina MiSeq platform was based on two steps of PCR. The first step amplified the mitochondrial 12S rRNA gene of amphibian species using the following primers: forward batra-F 5′-ACACTCTTTCCCTACACGACGCTCTTCCGATCTNNNNNNACACCGCCCGTCACCCT-3′

and reverse batra-R 5′-AGACGTGTGCTCTTCCGATCTNNNNNNGTAYACTTACCATGTTACGACTT-3′. Human blocking primer: TCACCCTCCTCAAGTATACTTCAAAGGCA-SPC3I was used to bind to human DNA and prevent its amplification (Valentini et al. 2016). This first PCR was performed in a final volume of 25 μL containing 5 μL of 5X Q5 reaction buffer, 0.2 μM “batra” primers, 0.2 mM dNTPs, 4 μM human blocking primer and 0.02 U/μL Q5 High-Fidelity DNA Polymerase (New England Biolabs) as well as 1 μL of positive control (mock community) or 5 μL of eDNA extract as a template. Amplification was performed in a Veritipro Thermal Cycler PCR Machine (Applied Biosystems, USA) under the following conditions: 30 s at 98°C, followed by 35 cycles of 30 s at 98°C, 30 s at 57°C and 1 min at 72°C and a final extension step at 72°C for 7 min (Valentini et al. 2016). Each sample was amplified in triplicate and then pooled before the purification step. We also used two negative PCR products to monitor contamination and the mock community as a positive control of the amplification in each PCR run. 1% agarose gel electrophoresis was run to check the successful amplification of the target band (170 bp) before purifying the PCR products using AMPure XP magnetic beads (Beckman Coulter, US).

Twenty forward and reverse index primers developed based on Sinclair et al. (2015) were used for Illumina sequencing. This second PCR was performed in a final volume of 20 μl containing 4 μl of 5X Q5 reaction buffer, 0.2 mM dNTPs, a combination of the indexed forward and reverse primers (0.25 μM each), 0.02 U/μl Q5 High-Fidelity DNA Polymerase (New England Biolabs) and 2 μL of purified first step PCR product. Amplification conditions were 30 s at 98°C, followed by 15 cycles of 10 s at 98°C, 30 s at 66°C, and 30 sec at 72°C, then a final extension step at 72°C for 2 min. The indexed target band was checked using 1% agarose gel electrophoresis. The second PCR product was purified using the same reagent as above, and quantified using the PicoGreen assay. The detailed DNA metabarcoding protocol for amphibians was published on protocols.io (<u>dx.doi.org/10.17504/protocols.io.973h9qn</u>). The first and second PCR mixtures were prepared in pre-PCR laboratory while the first purified PCR product was added in the post-PCR laboratory.

Equal amounts of DNA from each amplified sample were mixed to prepare the sequencing library. In addition to sequencing the 5 dilutions of the mock community, two or three environmental samples were spiked with one of the mock community species which was processed in the same sequence pool to confirm the presence and absence of the identified species. Furthermore, one random negative PCR product (entire sample) was included in each Illumina sequence run performed in 2019 and 2020. To detect all amphibian species in positive swab samples, the three swabs collected from positive Bd sites were sequenced while one swab was only sequenced from the other sites. The final pooled samples were visualized by gel electrophoresis to ensure that no extra unwanted fragments existed. The amplicon library was then sequenced on an Illumina MiSeq machine using the MiSeq v2 PE 150 Micro protocol, generating approximately 20 million paired-end reads of 150 bp length and exact matches to batra primers. Raw sequence data are available at NCBI through the SRA accession no. PRJNA821076 for 2019 and PRJNA822096 for 2020.

### Reference database construction

Amphibian 12S rRNA gene sequences were obtained by searching NCBI and BOLD using species names of the Norwegian amphibian species and ‘12S’ as search criteria. Next a homology search using Basic-Local-Alignment-Search-Tool (BLAST) was used to extract rRNA gene sequences of high (90%) homology. This two-step process led to a 12S rRNA gene sequence database of mostly amphibian sequences. To clean up the database, sequences were aligned and the “batra” primers were matched. This resulted in a database of 7 sequences including all species that have been reported in the wild in Norway.

### Observational data mining

Traditional observational data of amphibian diversity was obtained from the Norwegian public biodiversity databank “Artsdatabanken” (https://artskart.artsdatabanken.no/) . Data for amphibians was downloaded on 2020-05-11 using “Amphibia” as query resulting in 4086 species observations in the region overlapping with our samples. Species observations from the last 20 years were kept and then grouped if GPS coordinates overlapped as defined by resolution to the second decimal corresponding to a place up to 1.1 km^2^.

### Sequence analysis

Raw sequences were first processed with cutadapt v1.18 to remove PCR primers, and then analyzed with the R package dada2 v1.14.1 (Callahan et al. 2016) for denoising and sequence-pair assembly. After manual inspection of quality score plots, forward and reverse reads of the amplicon sequencing run were trimmed to 50 bp length. This assured an almost complete overlap of the forward and reverse reads as the amplicons without primers were 51-54 bp long. Additional quality-filtering steps removed any sequences with unassigned base pairs and reads with a single Phred score below 20. After reads were dereplicated, forward and reverse error models were created in dada2 with a subset of the sequences (10^8^ nucleotides). Amphibian 12S rRNA gene amplicons were assembled by merging the read pairs. Chimeras were removed using ‘removeBimeraDenovo’ from the dada2 package, which resulted in the final Amplicon Sequencing Variant (ASV) table for performing taxonomic assignments. Taxonomy (to species level) were assigned using blastn and an in-house 12S rRNA gene database. Sequences were assigned to specific species using cutoff criteria of identity > 96% (which corresponds to approximately two mismatches) and alignment length > 45 bp.

### Statistical analyses

Heatmaps were constructed using the ggplot2 R package, plotting the number of samples uniquely detected by either eDNA or traditional observation of species deposited to the Norwegian species database “Artsdatabanken” (ABD method), as well as those samples where the species was detected using both methods. Correspondence of species observations between eDNA and those recorded in Artsdatabanken (using GPS data rounded to the second decimal) were evaluated by correlation analysis, fisher exact test and Pearson’s Chi-square test using R (version 3.6.2). In order to trace the outcome of the sequence analysis we used the sequenced amphibian mock community.

Generalized linear (glm; base R) and additive (gam; “mgcv” package in R) models were used to explore detection rates of amphibian species in relation to time of sampling and pH. We also performed co-occurrence analysis using the R package “cooccur”. The later did not reveal any significant relationships probably due to sparse *Bd* detection.

## Results

### *Bd* detection and comparison of two TaqMan assays

We tracked the distribution of the invasive chytrid fungus *Bd* in the sampled water bodies. By using two TaqMan assays we were able to detect *Bd* in six water bodies and could confirm its presence in most cases by repeated sampling (Table 1). Still, our results indicate that spring is the season of highest detection probability for *Bd*, as samples taken during other seasons were all negative.

**Table 1.** List of positive *Bd* samples based on qPCR assay from 2017-2020.Vol: volume, Temp: temperature, Turb: Turbidity. Data points from 2017 are taken from Taugbøl et al. 2021, list of all samples with Bd results and amphibians species detected, are represented in supplementary Table S1 and S2.

| Sample ID | Date | Year | Vol | Temp | Turb | pH | Type | Locality | Assay |
| --- | --- | --- | --- | --- | --- | --- | --- | --- | --- |
| 10 | 26-May | 2017 | 600 | 15 | 0 | 7.32 | forest | Frogn | Taugbøl et al. 2021 |
| 13 | 26-May | 2017 | 180 | 15 | 3.61 | 6.71 | forest | Nesodden | Taugbøl et al. 2021 |
| 6 | 26-May | 2017 | 200 | 17 | 0.64 | 7.0 | agricultural | Ås | Taugbøl et al. 2021 |
| 3 | 26-May | 2017 | 190 | 15 | 31,83333 | 7.89 | agricultural | Østre Støkken | Taugbøl et al. 2021 |
| 1 | 26-May | 2019 | 540 | 12.5 | NA | NA | urban | Vårli, | Commercial |
| 3 | 26<br>May | 2019 | 19<br>0 | 15 | 31.8<br>3 | 7.8<br>9 | forest | Østre<br>Støkken | Commercial |
| 12 | 26-May | 2019 | 34<br>0 | 14 | 8.63 | 7.15 | urban | Ringveien<br>52 | Commercial |
| 69 | 01-Jun | 2020 | 60<br>0 | 24 | 2.16 | 7.9 | Agricult<br>ure | Østre<br>Støkken | Commercial &<br>Boyle |
| 75 | 01-Jun | 2020 | 36<br>0 | 23.4 | 4.47 | 7.09 | Suburb | Ringveien<br>52 | Commercial &<br>Boyle |
| 74 | 01-Jun | 2020 | N<br>A | 24.6 | 0.81 | 7.7 | Suburb | Odden | Commercial &<br>Boyle |
| Swab<br>38B | 30-May | 2020 | 54<br>0 | 21.9 | 6.33 | 7.8 | Agricult<br>ure | Værmyr,<br>Enningdal<br>en | Commercial &<br>Boyle |
| 10 | 25-May | 2020 | 48<br>0 | 18 | 2.12 | 5.96 | Forest | Butjenna | Boyle |
| 47 | 30-May | 2020 | 45<br>0 | 16 | 1.75 | 7.4 | Agricult<br>ure | Rokkevan<br>net | Boyle |

We used two TaqMan assays to amplify different regions of the rRNA operon (ITS1 and 5.8S rRNA gene). qPCR efficiency of the *Bd* 5.8S rRNA Genesig Advanced Kit and the Boyle TaqMan assay were on average 98.82% and 99.34%, respectively. The LOQ of the assays depending on the run varied between 100 and 10 gene copies for the commercial Genesig kit and 1 and 0.1 genome equivalents for the Boyle TaqMan assay (Figure 1; supplementary Table S3). The LOD as assessed from at least 3 independent runs varied between 10, 1 and 0.1 gene copies for the commercial Genesig kit and 1, 0.1 and 0.01 genome equivalents for the Boyle TaqMan assay (see supplementary Table S3). Thus it can be speculated that the Boyle TaqMan assay is slightly more sensitive than the Genesig kit assay. The higher sensitivity of the Boyle TaqMan assay was also confirmed using dilutions of the gBlocks fragment Bd_27-271 as standards, as the lowest concentration detected (in 2 of 3 triplicates) corresponds to 0.39 ITS1 copies (Figure 2B), while the LOQ (based on a single qPCR experiment) seems to be two orders of magnitude higher (39 ITS1 copies). It is well known that rRNA operon copy number per genome (or spore) is highly variable between strains of the same fungal species, as reported by Longo et al. (2013) for *Bd* strains genomes which contain from 10 to 144 ITS1 copies.

**Figure 1.**
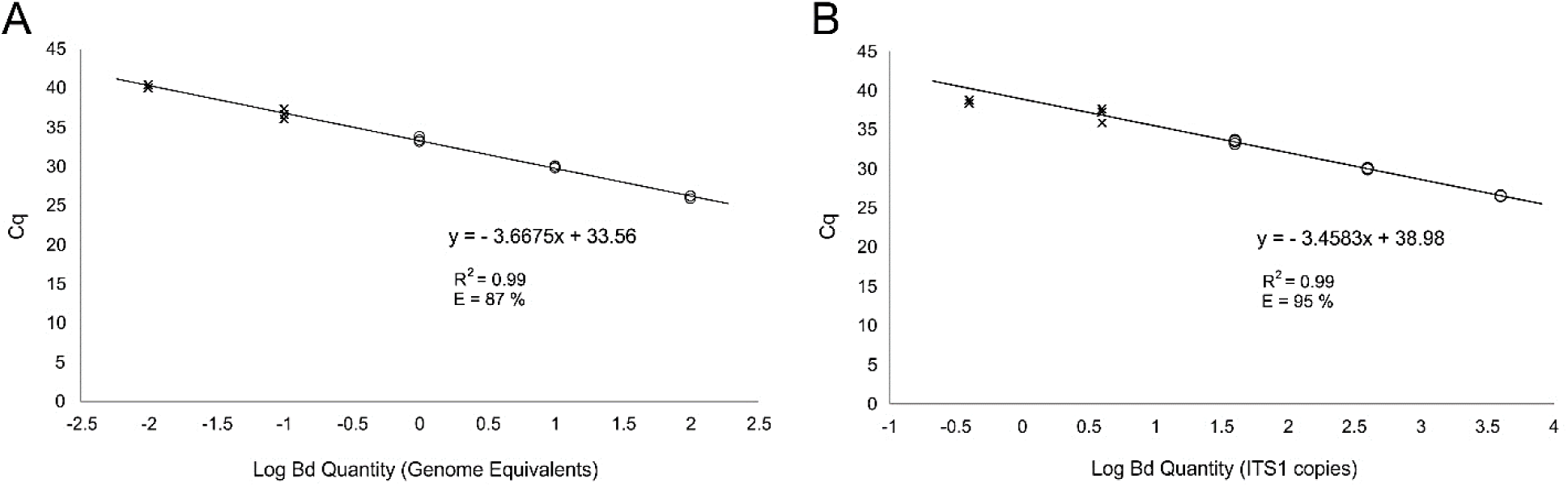
Examples for standard curves using to determine the limits of detection (LOD) and quantification (LOQ) for the Boyle TaqMan assay using spore DNA (providing genome equivalents) (A) and the synthetic gBlocks fragment *Bd*_27-271 (providing ITS copy numbers) (B) as standards. Crosses represent standard dilutions above the LOD but below the LOQ while circles represent standard dilutions used for determining the corresponding standard curves and their R2 and efficiency values (detailed on the panels). For the spore DNA curve (A), detailed statistics on multiple runs are provided in Supplementary Table S3.

**Figure 2.**
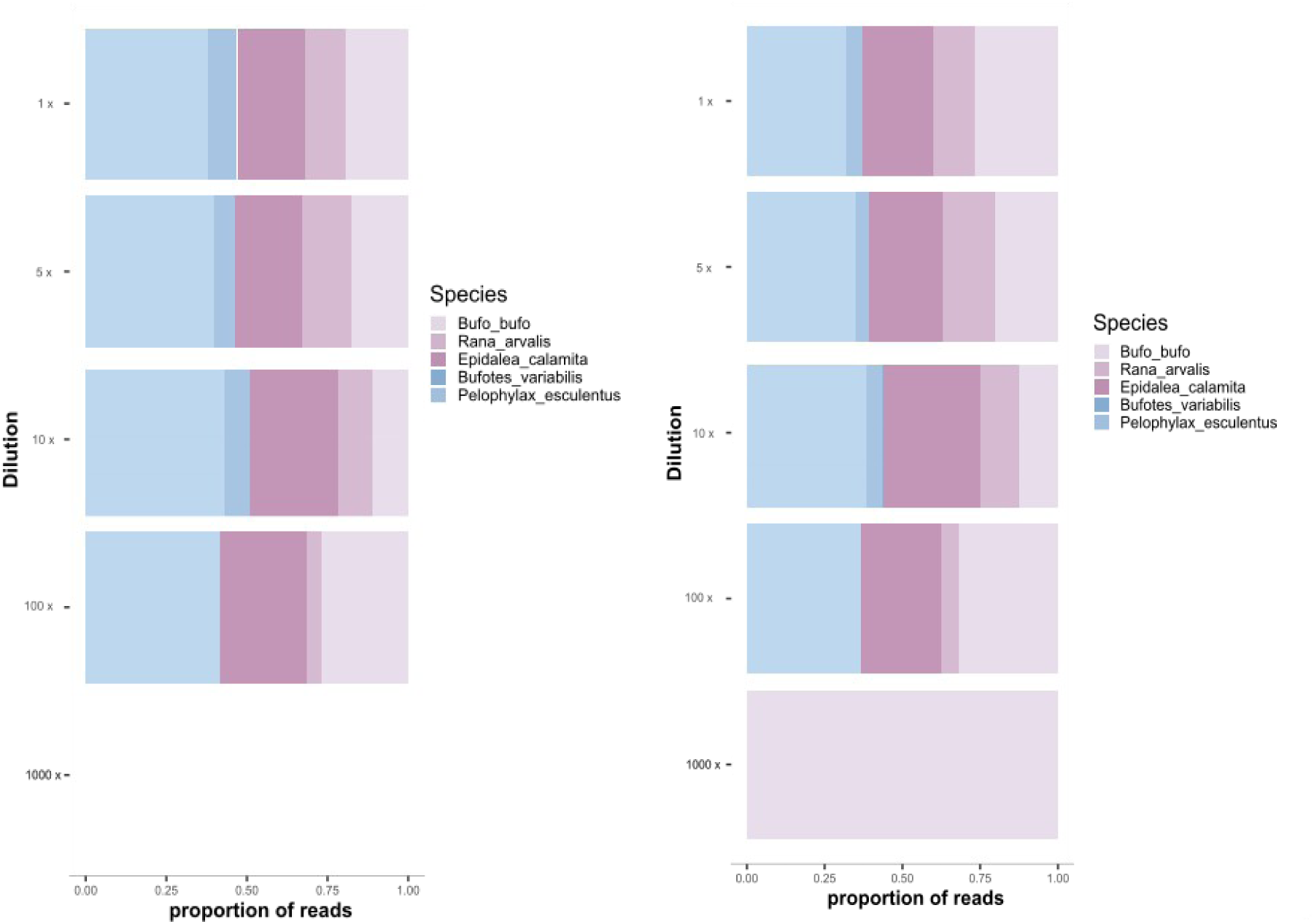
Sequencing results from the dilutions series of an amphibian mock community in 2019 (A) and 2020 (B). Both plots are showing the five added species in the original, 5 and 10 times diluted mock samples while in the 100 times diluted *Bufotes variabilis* disappeared.

Both qPCR assays generated positive *Bd* signals from water samples of three locations: Østre Støkken, Ringveien 52, and Odden, and from one swab from the location Værmyr (Table 1). Two additional locations, Butjenna and Rokkevannet, generated positive *Bd* signals when using the Boyle TaqMan assay. However, as a high Cq value (as in the case of Butjenna and Rokkevannet) can represent unspecific amplification and thus false positive *Bd* results, both PCR products were purified and Sanger-sequenced for validation. The resulting sequences were used to query the NCBI nt-database using blastn, which resulted in 100% matches with sequences from *Bd*, and thus presence of *Bd* was concluded.

### Internal positive control

We used internal positive controls (IPC) consisting of a fully synthetic DNA, primers and a TaqMan probe, which are included in qPCR reactions or coextracted with the DNA extract to detect PCR inhibition and false negative results due to inhibitory substances in the environmental samples (Hoorfar et al. 2004, Phillips, 2004). In 2019 and 2020, five and six samples, respectively, showed delayed Cq values above 27±2 for the IPC. This variability in Cq results strongly suggests the presence of inhibitory compounds in a minor fraction of samples, which needs to be taken into account when interpreting negative results (Supplementary Figure S2). The IPC was detected in all co-extracted samples indicating the efficiency of extraction protocol used. Three of these samples generated a normal IPC-qPCR signal after dilution, which is one way to overcome the effect of inhibitors. It should be mentioned that only half of these samples were successfully sequenced and one amphibian species was identified in each of them, thus metabarcoding results from these samples are likely biased as well. By reporting IPC-qPCR signal, potential inhibition can be observed providing a good indicator of false-negative PCR results. A way to solve some false-negative detection might be by sample dilution or optimizing qPCR conditions (Kamal et al. 2017, Lance and Guan, 2019) or the use of an inhibitor removal kit (McKee et al. 2015).

### Evaluation of amphibian metabarcoding

Amphibian diversity was assessed using a metabarcoding approach based on amplification of the mitochondrial 12S rRNA gene of amphibian species and subsequent sequencing. Positive controls such as the mock community including DNA from *R. arvalis, B. bufo, B. variabilis*, *E. calamita,* and *P. lessonae* showed amplification down to 10^-2^ ng of template DNA and resulted in sequences for most mock species (Figure 2). The exception was *B. variabilis* that could not be detected when the targeted DNA was below 10^-1^ ng of template DNA. Most of the species were detected in low concentration indicating that the metabarcoding approach is still sensitive to very low amounts of genomic DNA.

Next, we evaluated species discrimination of native Norwegian amphibian species by the DNA metabarcoding approach. The comparison of the available reference sequences by pairwise alignment revealed sufficient divergence amongst native Norwegian species for annotation at species level. The closest species were *R. arvalis* and *R. temporaria* with a minimum of two nucleotide differences in the amplified 12S rRNA gene region while all other species differed in at least four positions. Species discrimination may, however, become an issue with this metabarcoding approach in areas of high amphibian diversity including multiple closely related species such as in the tropics.

Analyses of the 110 and 91 samples from 2019 and 2020, respectively, resulted in all of the samples being well amplified as reflected by the amplified amplicon size (approx. 50 bp, expected to the target region). No amplification was obtained from negative control filters, thus allowing us to exclude the possibility of cross-contamination among filters and confirming the clean DNA extraction and purification steps. Moreover, there was no amphibian species detected in any negative PCR control of the metabarcoding after the end-point (no resulting sequences), even if a band appeared in gel electrophoresis due to the primer dimer formation, confirming a clean PCR setup.

A stringent raw sequence data quality filtering resulted in a loss of approximately 60% of the reads from an average of 15,736 raw paired end reads per sample (range from 830 to 87,543). From an average of 6,181 (range from 587 to 29,720) quality-filtered paired end reads per sample, denoising and merging resulted in a mean number of 5,676 high quality sequences (range from 567 to 29,136). Only in 68 of 110 samples collected in 2019, and in 69 of 91 samples from 2020, amphibian sequences could be detected. Unspecific amplification of non-targeted species could be observed in almost all amplified samples. In 2019 and 2020 the mean percentages of sequences per sample assigned to amphibian taxa were 9% (range 0 to 96%) and 30% (range 0 to 100%), respectively, corresponding to 947 (range 0 to 26,327) and 1,862 (range 0 to 12,667) amphibian reads per sample. Closer inspection of the non-amphibian sequences showed various taxonomic assignments from bacteria to fish. This indicates unspecific amplification in the PCRs and emphasizes the need for a high sequencing depth to detect amphibian species when using the batra primers. Still, as we obtained on average more than 5,000 paired reads per sample, false negatives as a result of sequence library preparation could be minimized.

Looking at 2019 data, amphibian detection was significantly lower in September compared to May/June and July (estimate = −1.635, z-value = −3.710, p-value = 0.0021 as given by generalized linear models). This was corroborated by the total number of detected amphibian occurrences among all sites (Table 2) and at six sites that were sampled during the three sampling occasions in 2019. In autumn amphibians could not be detected at any of the six sites while in May/June and July there were multiple species detections. Thus it can be concluded that sampling during spawning and post-spawning results in higher likelihood of detection when compared to autumn sampling. In a previous study, a comparison of the detection of the pool frog (*P. lessonae*) using both eDNA and traditional methods revealed that traditional methods gave a higher rate of observation in June, whereas eDNA gave at least as many or more observations during other months (Eiler et al. 2018).

**Table 2.** Comparison of species detection among the three sampling times (May, July and September) in 2019. Numbers represent the number of sites where a species was detected, clearly showing that the number of amphibian species positive sites was lowest in September. List of all samples, raw sequences, sampling volume and environmental parameters are represented in supplementary Table S1 and S2.

| species | May/June | July | September |
| --- | --- | --- | --- |
| *Rana arvalis* | 2 | ND | ND |
| *Rana temporaria* | 9 | 2 | 3 |
| *Bufo bufo* | 10 | 3 | 1 |
| *Triturus cristatus* | 6 | 10 | ND |
| *Lissotriton vulgaris* | 9 | 14 | 3 |

### Amphibian species distribution in Norway

Five amphibian species *B. bufo*, *L. vulgaris*, *R. arvalis*, *R. temporaria*, and *T. cristatus* are prevalent in Norwegian waterbodies. This limited number of species was observed by our eDNA approach with more than one amphibian species being detected in several samples. A study on the influence of pH on amphibian diversity in Norway reported that *R. temporaria* was detected in all pH levels but there is a decrease in the frequency of *B. bufo*, *T. vulgaris* (Dolmen et al. 2009), while *R. arvalis* and *T. cristasus* seemed to be sensitive to pH changes. Using our eDNA data and generalized additive models revealed a significant decreases for *R. arvalis* (Estimate = −1.388, z-value = −2.958, p-value = 0.0031) and *T. cristatus* (Estimate = −3.566, z-value = −3.059, p-value = 0.0023) stronger than for *L. vulgaris* (Estimate = −0.979, z-value = −2.186, p-value = 0.0288) and *R. temporaria* (Estimate = −0.789, z-value = −2.013, p-value = 0.0441). Furthermore, our results confirm previous conclusions that acidic areas are lower in amphibian diversity in the inland/highland region of Southern Norway as given by generalized additive models revealing a significant decrease in overall species observations with decreasing pH (Estimate = −3.350, z-value = −3.644, p-value = 0.0003).

### Comparison between eDNA and traditional observation monitoring methods

Five amphibian species were detected by both traditional observation methods (ABD) and eDNA (Figure 3). A much larger number of sites has been covered by ABD when compared to eDNA methods. Still, there were 133 sites where overlapping data was available for comparative analysis. An all data heatmap is shown in Figure 3. Fisher exact and Pearson’s Chi-squared tests indicate a significant association between eDNA and ABD in detecting four of the amphibian species while no significant association could be observed for *R. temporaria* (Table 3). To conclude, correspondence of species observations between data from a national species reporting ABD-based program and eDNA metabarcoding was significant but weak. This may reflect both an incomplete overlap in species detection and differences in the number of total detections of individual species, and the different timeframes (20 years vs 2 years).

**Figure 3.**
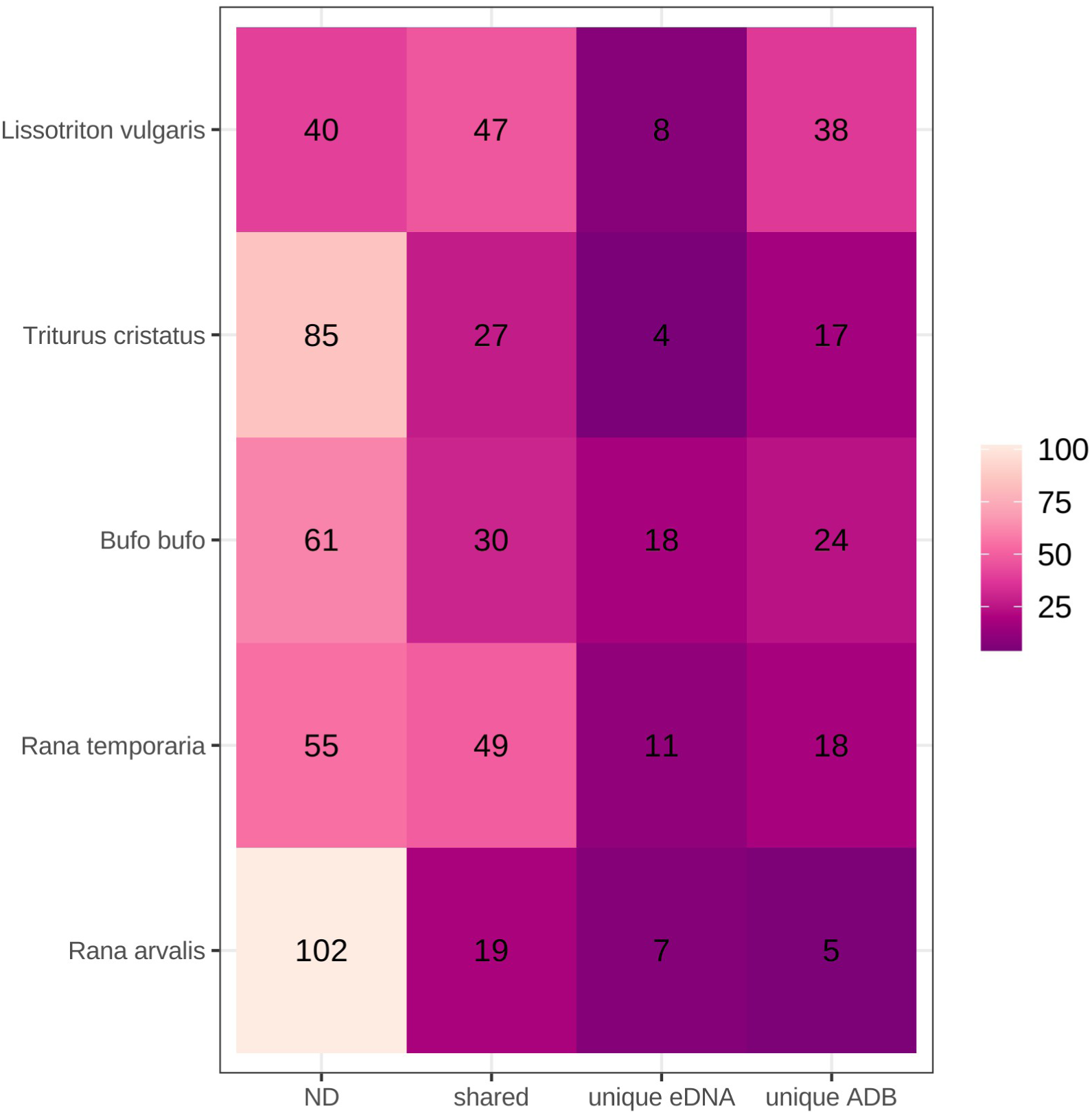
Comparison of amphibian species detection between our metabarcoding results (2 years of data) and species inventories in the Norwegian species database “Artsdatabanken” (20 years of data). ND – number of sites where the respective species was not detected by both methods; shared – the number of sites where the respective species was detected by both methods; unique eDNA – the number of sites where the respective species was detected only by eDNA; unique ABD - the number of sites where the respective species was found only in the database. Detailed statistics on correspondence between methods are given in Table 3.

**Table 3.** Detailed statistics on the comparison of amphibian species detection between our metabarcoding results and species inventories in the Norwegian species database “Artsdatabanken”, whose results are shown in Figure 3.

|  | Correlation | Fisher exact test (odds | Pearson's Chi-squared |
| --- | --- | --- | --- |
| Species | (R/p value) | ratio/p-value) | test (p-value) |
| *Rana arvalis* | 0.24/<0.001 | 5.37/0.017 | 0.016 |
| *Rana temporaria* | 0.13/<0.001 | 2.17/0.204 | 0.2 |
| *Bufo bufo* | 0.25/<0.001 | 3.70/0.006 | 0.004 |
| *Triturus cristatus* | 0.41/<0.001 | inf/<0.001 | <0.001 |
| *Lissotriton vulgaris* | 0.21/0.04 | 3.56/0.017 | 0.02 |

## DISCUSSION

A major advantage of metabarcoding is that they facilitate the analysis of multiple taxonomic groups from the same sample. In principle, biodiversity across the entire tree of life present in a particular system can be assessed by DNA metabarcoding (Stat et al. 2017). In addition to amphibian diversity, we tracked the distribution of the invasive chytrid fungus *Bd* in the sampled water bodies.

Similar studies have found *Bd* in various water bodies from other Nordic countries in Denmark and Sweden, and also in association with infected amphibians using genetic assays (Rosquist 2020). In this previous study, it was argued that skin swabs provided a higher likelihood of *Bd* detection. However, the comparison of detection probability between the two methods in the Rosquist (2020) study was highly biased, as only a single water sample was taken from an individual system, while swabs were taken from multiple specimens (up to 97) during multiple sampling occasions spread over several years from a single system. We therefore attempted to assess detection probabilities of the two methods using comparable sampling efforts, i.e. using the same amount of time to take water and swab samples. Since it takes a much longer time to obtain swab sample than taking single water sample, we only obtained very few swab samples. Not only time-efficiency but also with hindsight of non-destructive water sampling should be the preferred method. However, the low number of *Bd* positive waterbodies in Norway did not allow us to perform a thorough comparison. Still, confirming results with diverse methods such as swab-and water-based detection, as well as multiple molecular assays, is recommended since this can increase detection probability and confidence in positive detection.

### Lessons Learned for a national monitoring program based on eDNA

Screening amphibian occurrence by field techniques usually requires a suite of tools, which introduce several sources of errors. One of the errors is the high variability of amphibian detection at different seasons throughout the year (Heyer et al. 1993) and another is error in the sampling technique. For example different types of amphibian traps affect detection probabilities (Dodd, 2010; Petitot et al. 2014). This has been proposed by previous observations such as in neotropical streams in Brazil where both ABD and eDNA metabarcoding methods were used (Sasso et al. 2017). Our results confirm previous observation that amphibian should be monitored in late spring early summer in Scandinavia when using eDNA methods (Eiler et al. 2016). Thus, national monitoring programs for amphibians based on eDNA are best performed from late May to early July.

The high dispersion and low detection probability represents a challenge for monitoring programs in risk assessment, as large areas including high numbers of systems need to be monitored. There is also a challenge to integrate the monitoring results in management strategies to protect amphibian diversity. To map large areas, such as across Norway, a random sampling strategy should be set up. In our case for the risk assessment of Bd, we focused on systems impacted by humans since the main pathway of introduction of Bd is related to human activities (VKM report). Since sampling of a vast amount of systems is necessary efficient sampling techniques with low likelihood of contamination need to be applied. We chose a low tech approach using a sterile beaker attached to expandable (easy to clean) sampling pole, sterile syringes and cartridge filters. This kept the time spent per site to a minimum (20-45 minutes) including the assessment of water parameters with online sondes.

For sample preservation, we recommend flash freezing of filters in liquid nitrogen on site using a dry shipper and transfer to −80 freezers in the lab as this allows the long-term preservation of all kinds of environmental and tissue samples. This has been the “golden standard” for downstream DNA and RNA analysis for at least three decades (Frampton et al. 2008, Kilpatrick et al. 2002, Seutin et al. 1991, Reiss et al. 2012, Wong et al. 2012, Anchordoquy and Molina, 2007). For DNA extraction, we recommend commercial kits that are optimized for filter cartridges and remove potential PCR inhibitors while still keeping DNA yield high, i.e. the DNeasy PowerWater Sterivex kit. This is an important initial step of the analysis workflow that needs to be kept constant as extraction protocols determine to a large extent the outcome of metabarcoding studies (Majaneva et al. 2018). For library preparation in the case of metabarcoding we strongly argue for a two-step PCR. There are several reasons: 1) Short adapters with undefined nucleotides between target primer and sequencing primer minimize biased PCR amplification in the first PCR, 2) a two-step PCR minimizes heteroduplex (chimera) formation (Polz et al. 2003). 3) Indexing in second PCR which adds also remaining adapter sequence prevents index jumping.

Bioinformatic workflows for national monitoring programs should use denoising of reads based on quality scores and optimized for each sequencing run as for example implemented in the dada2 algorithm (Challahan et al. 2016) instead of clustering to improve species resolution. Using such methods allows discrimination of species with 2 nucleotide mismatches. Reference databases should be developed specific for the national monitoring project taken from national species inventories including potential invasive species. This was rather simple in our case of seven amphibian species being present in Norway, and might be more challenging in species rich taxonomic groups or locations. Species resolutions of the metabarcoding assay can be tested in silico using for example database-against-database homology searches and lowest common ancestor analysis (Huson et al. 2007).

In the case of qPCR we recommend the use of IPC to identify samples that exhibit PCR inhibition. These samples can be flagged as problematic and run again after dilution which can lead to positive detection of targets. There is also a need to decrease LOD and LOQ keeping amplification efficiency between 90-110 %. The lowest number of replicate PCR reactions should be three and if budget constrains allow can be increased (Piggott et al, 2016). In addition, high Ct samples should always be evaluated by sequencing to detect false positives.

Besides these recommendation there are still many obstacles standing in the way for the implementation of eDNA based biodiversity monitoring. International coordination is paramount to ensure comparability of data in time and space and to avoid unnecessary duplication of efforts and resulting delays in wide application. Awareness and knowledge of the new methods among practitioners is essential for implementation and to provide platforms for critical discussion on when, where and how the eDNA based methods should be applied. Standardization efforts need to advance and cover all steps from sampling design to sequence data interpretation. Additionally, in order to allow for comprehensive mapping of genetic diversity, the number of published genomes (at least mitochondrial genomes) should cover all known species. Modelling and analysis tools should be applied and developed alongside to facilitate efficient decision making.

### Putting our results into context of previous risk assessment for amphibians due to Bd

Our study shows that eDNA surveys provide detailed insights into the distribution of amphibians and an associated pathogen, thus allowing the assessment of risks associated with the invasive chytrid fungus *Bd*. Reproducible detection of *Bd* over multiple years strongly suggests that this pathogen is established at multiple sites in Southern Norway. *Bd* positive samples were obtained throughout an area of at least 1000 km^2^ south of Oslo with *Bd* detection highly dispersed within the area, which is comparable to similar climatic regions in Sweden and UK.

Multiple methods have been proposed to assess risks for biodiversity including Species Distribution Models (SDMs) combined with species specific biotic indices (Rödder et al. 2009), expert opinion (Non-native Risk Assessment scheme; Roy et al. 2013) and an integrative analytical approach combining ecological traits and anthropogenic risk factors (Munstermann et al. 2021). *Bd* is known to be a particularly temperature and moisture dependent species (Lips et al. 2008, Woodhams et al. 2008). The *in vitro* growth optimum range of *B d* is at 17–25 °C, whereas temperatures higher than 29 °C, freezing and desiccation are lethal (Piotrowski et al. 2004). These findings are supported by observations in the field (Kriger et al. 2007). A previous study on the geographic extent of *Bd*’s climatic niche based on SDM suggested presence but low risks of *Bd* for anuran amphibian species in Norway (Rödder et al. 2009). In another recent study it was argued that transmission from an environmental *Bd* reservoir and climate change could increase the ability of *Bd* to invade new amphibian populations and increase their extinction risk. Still, a recent study concludes that *Bd*-induced extinction dynamics were far more sensitive to host resistance and tolerance than to *Bd* transmission (Wilber et al. 2017). Infection-tolerant host species seem to dominate in Norway, as suggested by previous European studies (Smith 2014). Hence, the overall likelihood of a *Bd* negative impact on indigenous amphibian species has been suggested to be with moderate confidence minimal while for an introduced frog species (*P. lessonae*) there was a moderate risk identified (VKM, et al. 2019). Considering previous outbreaks of chytridiomycosis in Europe (Bosch and Martinez-Solano 2006) and extensive studies on *Bd* in similar climate regions, as well as its low prevalence in our study coupled with co-occurrences of various indigenous species over multiple years, it can be assumed that chytridomycosis outbreaks in Norway are unlikely. Still, it can be argued for annual follow up of *Bd* presence at least for the positive sites and nearby areas to keep an eye on the spread of this fungal pathogen.

## Acknowledgement

We would like to thank Mats Töpel and Thomas Larsson for their help in planning this work. Funding was provided by the Norwegian Environment Agency.

---

**Supplementary Table S1** List of samples from 2019 with raw sequences, sampling volume and environmental parameters.. Samples with positive (+) and negative (-) *Bd* results and amphibian species detected are represented.

**Supplementary Table S2** List of samples from 2020 with raw sequences, sampling volume and environmental parameters. Samples with positive (+) and negative (-) *Bd* results and amphibian species detected are represented. All swabs collected from positive *Bd* sites were sequenced while one swab was only sequenced from the negative *Bd* sites.

**Supplementary Table S3.** Summary statistics on the limit of detection (LOD) and the limit of quantification (LOQ) of the *Bd* assays. LOD and LOQ: 10, and 1 gene copies for commercial kit and 1 and 0.1 genome equivalent for Boyle TaqMan assay.

**Supplementary Figure S1.** Sampling scheme of the sampling in 2019 showing the distribution of sampling locations during the three sampling occasions (A). Distribution of *Bd* as assessed by the analysis of water samples, swabs, and tadpole DNA around Oslo (B) and Southern Norway and Central Sweden (C). In the latter map the distribution of all samples obtained in this study (data from 2019 and 2020) are shown. Additional data was obtained from Rosquist (2020) and Taugbøl et al. (2021). Full symbols indicate detection of *Bd* by the TaqMan assays while open symbols indicate negative results.

**Supplementary Figure S2.** Scatter plot of Cq values of standard and samples multiplexed with the internal extraction control. The figure only shows 3 samples with Cq values more than 27 and less than 50 while the other 3 samples had Cq value more than 50, not detected by qPCR.

## Notes

### Competing Interest Statement

The authors have declared no competing interest.

