## Supplementary Material for "Seasonal eDNA-based monitoring of *Batrachochytrium dendrobatidis* and amphibian species in Norway"

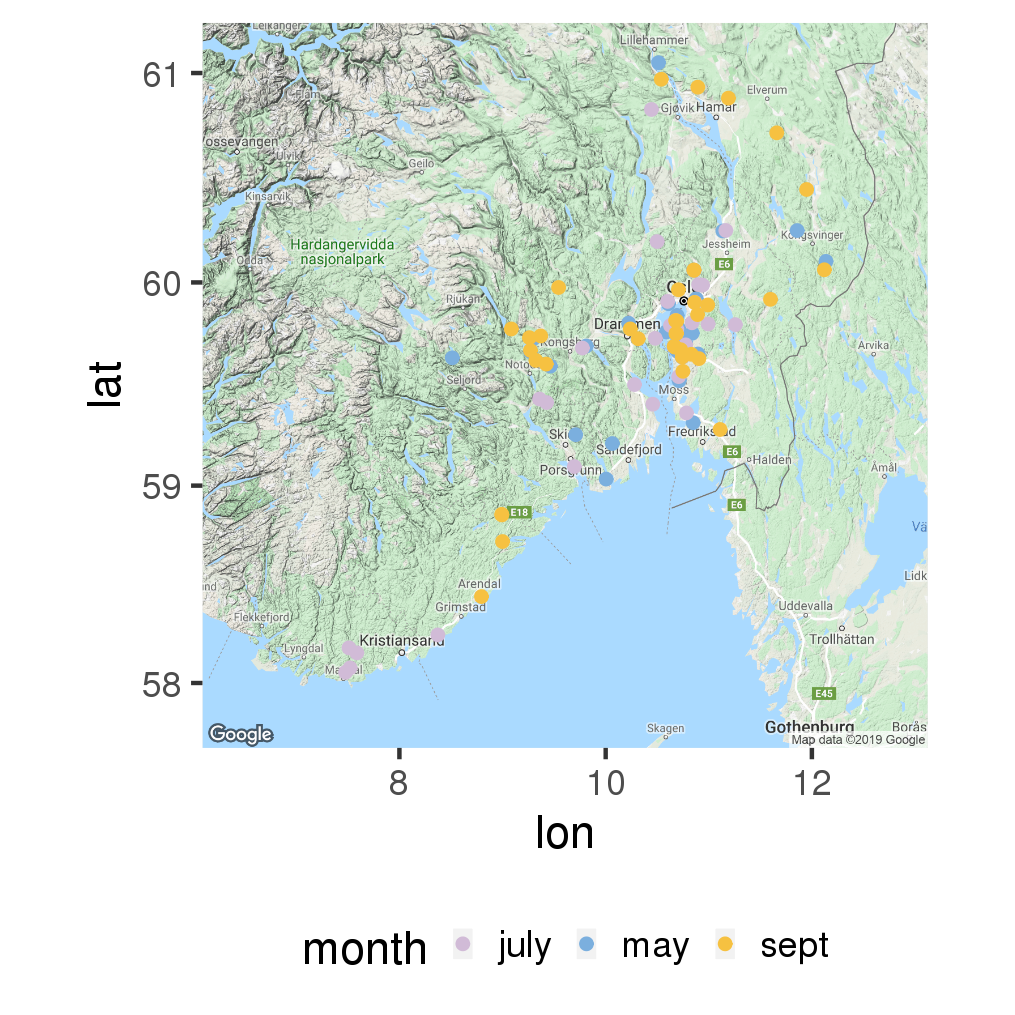
**Supplementary Figure S1A.** Sampling scheme of the sampling in 2019 showing the distribution of sampling locations during the three sampling occasions.

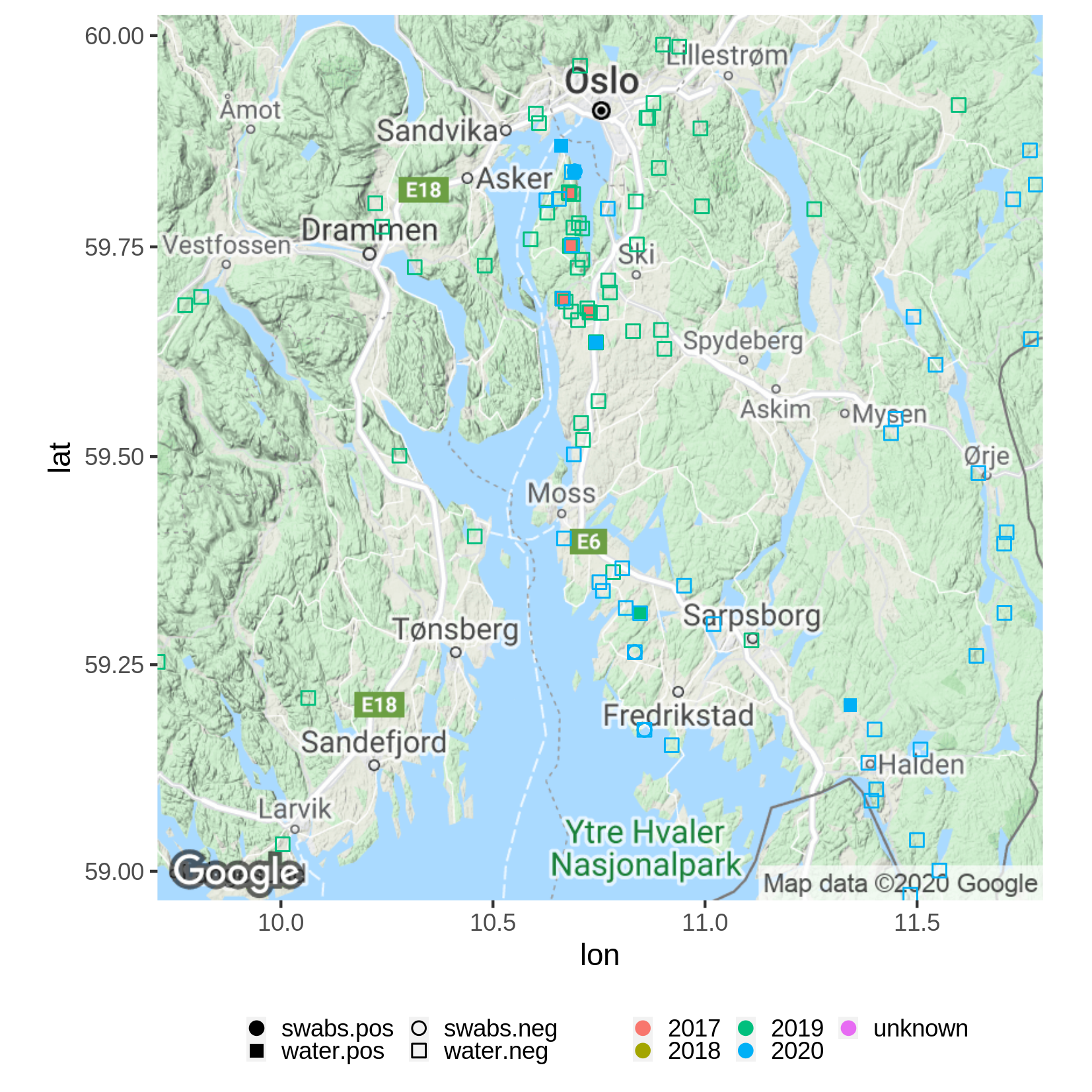

**Supplementary Figure S1B.** Distribution of *Bd* as assessed by the analysis of water samples, swabs, and tadpole DNA around Oslo

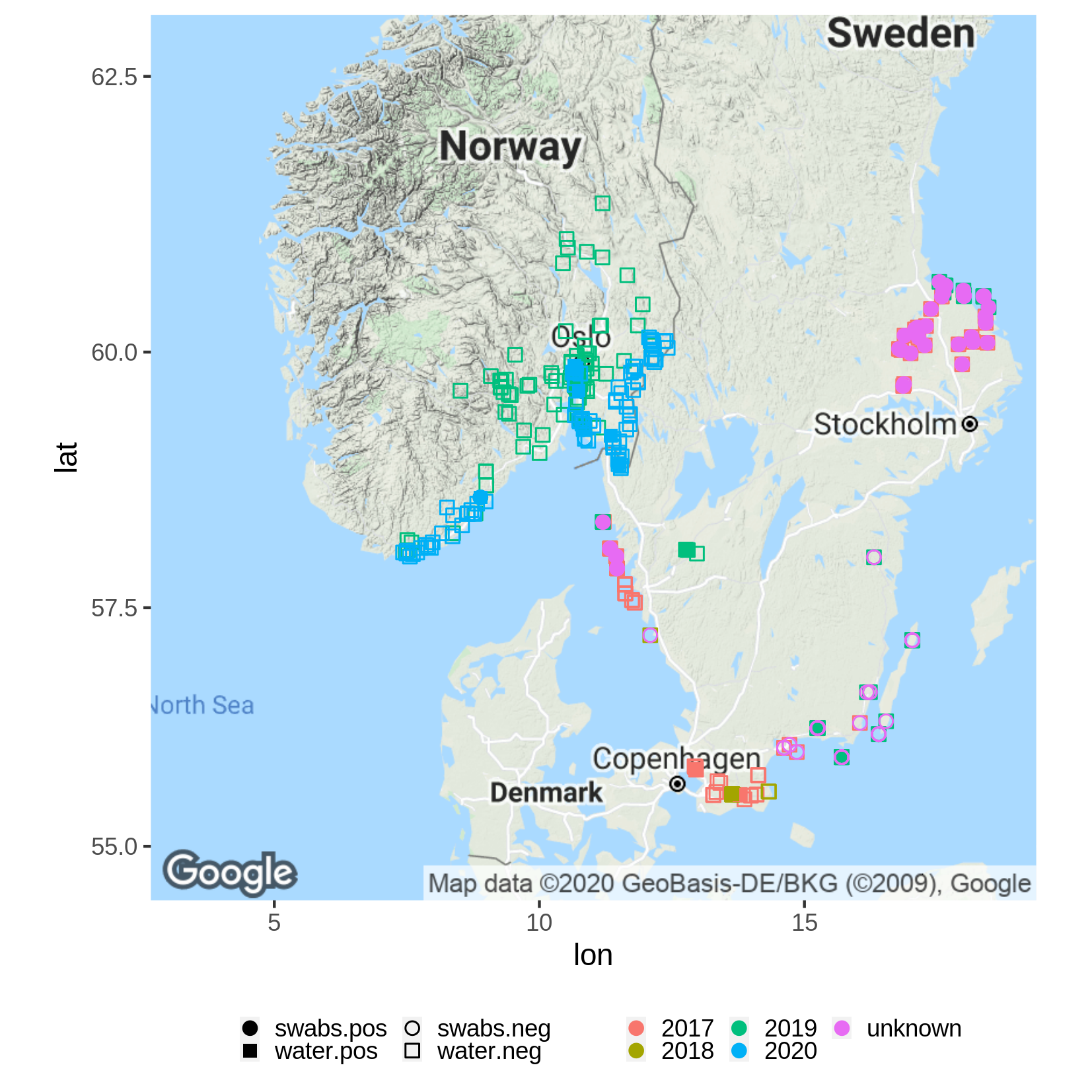

**Supplementary Figure S1C.** Distribution of *Bd* in Southern Norway and Central Sweden including all samples obtained in this study (data from 2019 and 2020) are shown. Additional data was obtained from Rosquist (2020) and Taugbøl et al. (2021). Full symbols indicate detection of *Bd* by the TaqMan assays while open symbols indicate negative results.

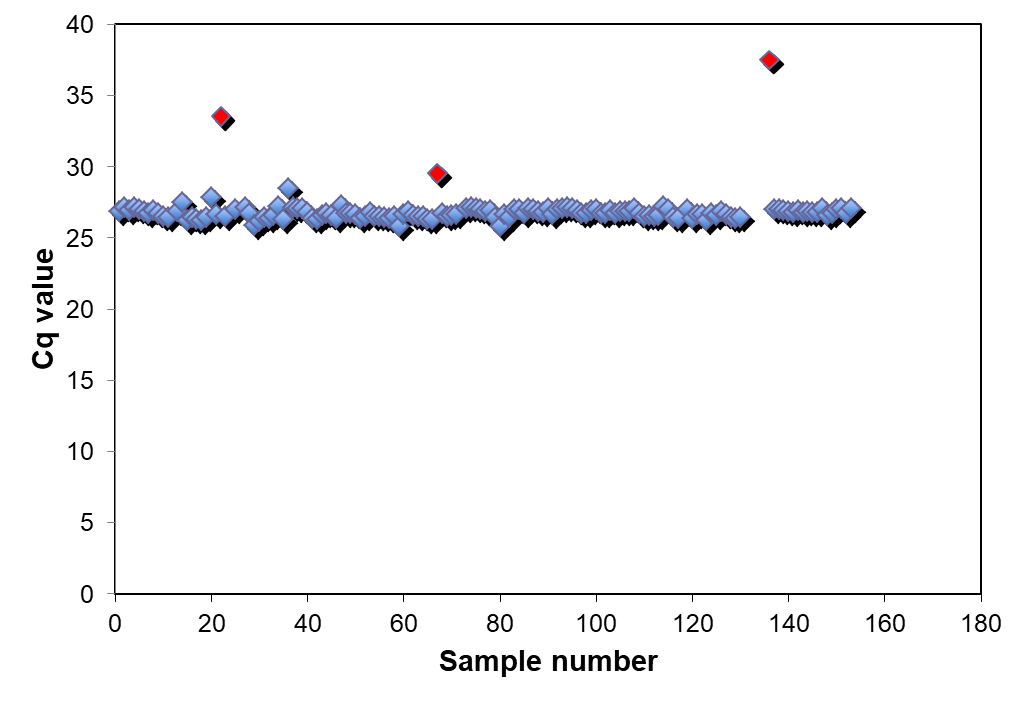

**Supplementary Figure S2**. Scatter plot of Cq values of standard and samples multiplexed with the positive internal control from year 2020. The figure only shows 3 samples with Cq values more than 25 and less than 50 (red color) while the other 3 samples had Cq value more than 50, not detected by qPCR.

**Supplementary Table S3**. Summary statistics on limit of detection (LOD) and limit of quantification (LOQ) of the *Bd* assays. LOD and LOQ: 100 and 10 gene copies for commercial kit and 1 and 0.1 genome equivalent for Boyle TaqMan assay.

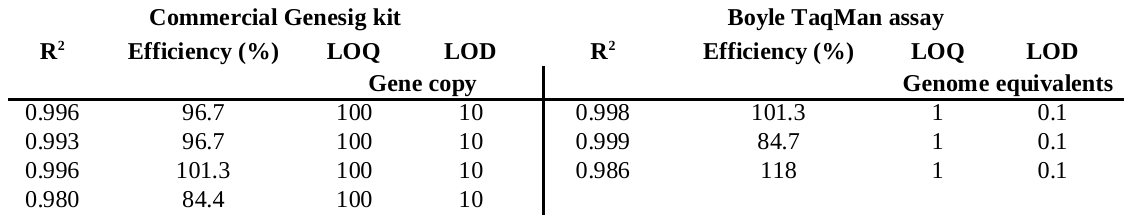
